# *PhenoImage*: an open-source GUI for plant image analysis

**DOI:** 10.1101/2020.09.01.278234

**Authors:** Feiyu Zhu, Manny Saluja, Jaspinder Singh, Puneet Paul, Scott E. Sattler, Paul Staswick, Harkamal Walia, Hongfeng Yu

**Affiliations:** Department of Computer Science and Engineering, University of Nebraska-Lincoln, USA; Department of Agronomy and Horticulture, University of Nebraska-Lincoln, USA; Wheat, Sorghum and Forage Research Unit, USDA-ARS, Lincoln, NE, USA

**Keywords:** high-throughput phenotyping, image processing, plant phenotyping, RGB images, fluorescence images, sorghum, wheat

## Abstract

High-throughput genotyping coupled with molecular breeding approaches has dramatically accelerated crop improvement programs. More recently, improved plant phenotyping methods have led to a shift from manual measurements to automated platforms with increased scalability and resolution. Considerable effort has also gone into the development of large-scale downstream processing of the imaging datasets derived from high-throughput phenotyping (HTP) platforms. However, most available tools require some programing skills. We developed *PhenoImage* – an open-source GUI based cross-platform solution for HTP image processing with the aim to make image analysis accessible to users with either little or no programming skills. The open-source nature provides the possibility to extend its usability to meet user-specific requirements. The availability of multiple functions and filtering parameters provides flexibility to analyze images from a wide variety of plant species and platforms. *PhenoImage* can be run on a personal computer as well as on high-performance computing clusters. To test the efficacy of the application, we analyzed the LemnaTec Imaging system derived RGB and fluorescence shoot images from two plant species: sorghum and wheat differing in their physical attributes. In the study, we discuss the development, implementation, and working of the *PhenoImage*.

**Highlight:** *PhenoImage* is an open-source application designed for analyzing images derived from high-throughput phenotyping.

## Introduction

In the genomics and post-genomics era, technological advances in sequencing platforms have paved the way for high throughput genotyping (Jackson *et al*., 2011; Furbank and Tester, 2011). These developments coupled with molecular breeding approaches have enhanced the genetic understanding of plants, which has dramatically progressed the crop-improvement efforts (Moose and Mumm, 2008; Varshney *et al*., 2009; Tester and Langridge, 2010). However, precise and efficient phenotyping has been a challenge (Furbank and Tester, 2011). To tackle this problem, plant phenotyping technologies have achieved a huge leap in recent times; the shift from laborious and error-prone manual measurements towards automation (Fiorani and Schurr, 2013; Gong and He, 2014). Automated imaging-based platforms have tremendously enhanced our ability to record a plant’s physical and physiological attributes in a non-invasive manner. Despite these advances, phenotyping technologies still trail developments on the genomics front, especially the rate at which the phenotypic data is generated (Houle *et al*., 2010; Furbank and Tester, 2011; Minervini *et al*., 2015; Gehan *et al*., 2017). The major limit is not the ever-evolving sophisticated instrumentation for image capturing but with the downstream processing of large-scale phenotypic data, which is not easily accessible to many plant biologists.

High-throughput (HTP) imaging platform refers to the accurate acquisition and analysis of multidimensional traits at the individual plant level in context of this work (Yang *et al*., 2020). To no surprise, these platforms generate a diversity of images corresponding to different spectra of light such as RGB, near-infrared, fluorescence, and hyperspectral. Thus, terabytes of digital information can be routinely generated through an imaging experiment. Currently, the website: www.plant-image-analysis.org, documents 179 image software tools (Lobet *et al*., 2013). Availability and usage of some of these tools are being restricted and adheres to proprietary rights, for instance LemnaGrid-Scanalyzer3D by LemnaTec GmbH, Germany. On the other hand, several open-source tools designed for specific applications, ranging from cell to whole canopy analysis, are readily accessible in the public domain (Lobet *et al*., 2013). In addition to their broader functionalities these have also opened new avenues to integrate third-party algorithms. Examples include HTPheno (developed as a plugin for ImageJ) (Hartmann *et al*., 2011), Plant Computer Vision or Plant CV (a community-based toolkit for plant phenotyping analysis) (Fahlgren *et al*., 2015; Gehan *et al*., 2017), Integrated Image Platform or IAP (Klukas *et al*., 2014), Image Harvest (Knecht *et al*., 2016). Despite their power and flexibility, these tools may require some proficiency with programing language as a pre-requisite to process large-scale datasets. This is a challenge for many biologists with limited or no coding skills.

The availability of several affordable automated and semi-automated phenotyping platforms has increased their usage to score the traits of interest (Klukas *et al*., 2014; Li *et al*., 2014). Keeping this view in mind, we developed *PhenoImage* – an open-source, GUI-based cross-platform solution for large-scale data processing that is not only convenient to use but highly precise and effective at the same time. The intuitive nature of the application will allow plant scientists with little or no knowledge of programming language to process phenotypic dataset on their personal computers. In addition, the application can facilitate parallel processing of large-scale image data on high-performance computing clusters. To test the efficacy of the application, we analyzed the LemnaTec Imaging system-derived RGB and fluorescence images from sorghum and wheat, which differ in their physical attributes. The availability of multiple functions and filtering parameters provides flexibility to analyze a wide variety of plant species. Images acquired from other phenotyping platforms or handheld devices can also be processed using *PhenoImage*.

## Materials and Methods

### *PhenoImage* Workflow

*PhenoImage* is a MATLAB-based application i.e. compatible with multiple operating systems. The software is available in two versions, a regular version that requires MATLAB license and a standalone version that uses ‘MATLAB Compiler Runtime’ and does not necessarily require MATLAB license for its operation. Both versions can be downloaded from: http://wrchr.org/phenolib/phenoimage. The same graphical user interface (GUI) application can support image data processing on a single central processing unit (CPU) as well as parallel processing on High Performance Computing (HPC) clusters.

### Software Development and Implementation

The GUI for high throughput image analysis is based on MATLAB. The summary of image processing workflow of *PhenoImage* includes: (i) file loading, (ii and iii) image cropping and filtering, (iv) digital trait extraction using specific functions (based on the user’s requirement), (v) followed by image processing either on a local machine or HPC clusters (Fig. 1). We have provided a step-by-step guide to use *PhenoImage* (see *PhenoImage* Guide Document).

**Fig. 1.**
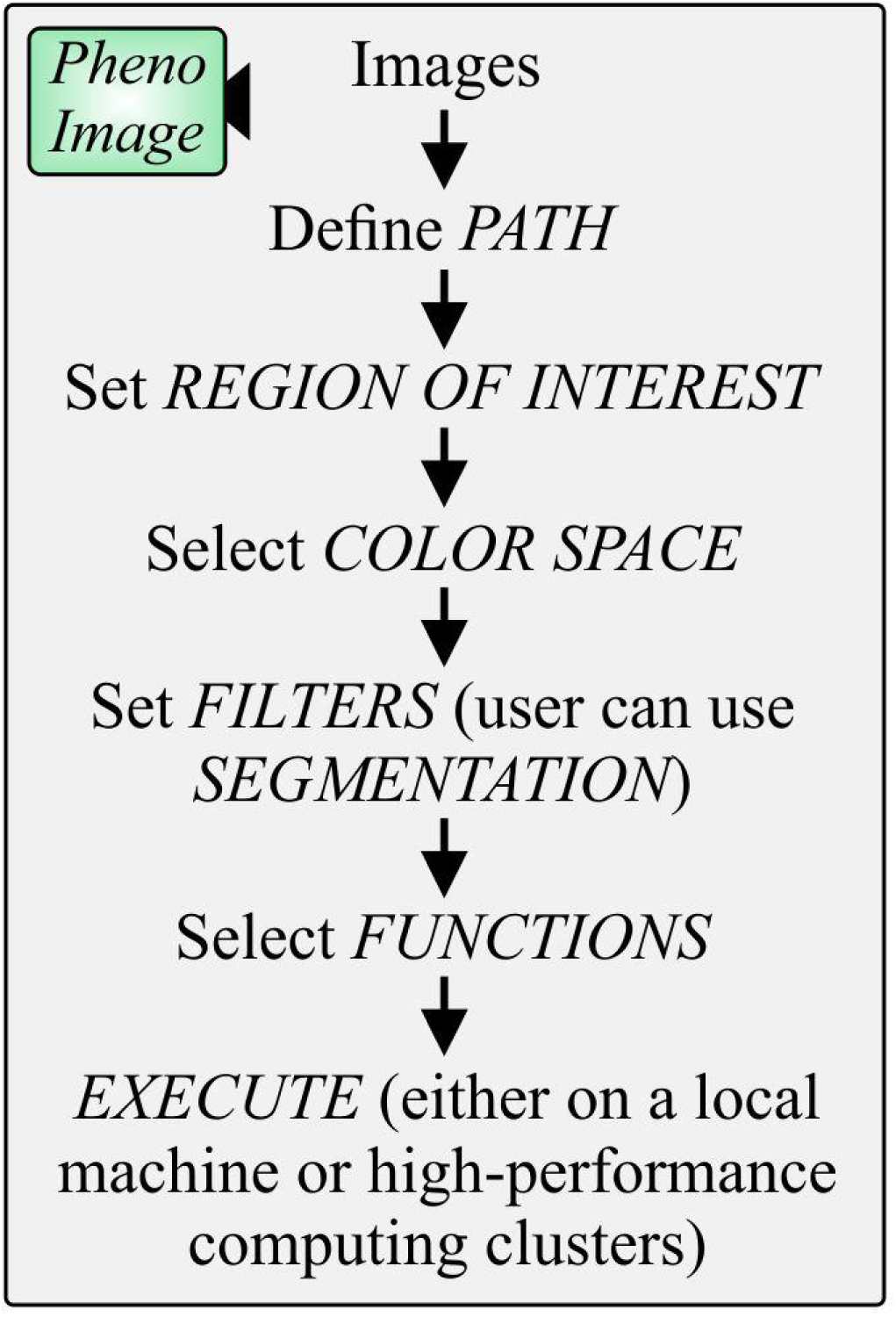
*PhenoImage* Workflow. First, the path of the folder containing RGB images is described. Then, a region of interest containing plant image is defined, followed by the selection of color space of preference. Afterwards, filter parameters are determined, and functions corresponding to the digital traits of users’ choice are selected. Then, the processing of the images is executed using either a local CPU or high-performance computing clusters.

### File loading

The image files can be loaded by specifying regular expressions in ‘*Path*’ using the following format: “FOLDER NAME*\\**.*png*”. This should allow the loading of all images under the respective folder. The application is compatible with widely used image formats such as *jpg, png*, and *tiff*. The spinner can be used to change the ‘*Original Image*’ that is currently displayed (Fig. 2).

**Fig. 2.**
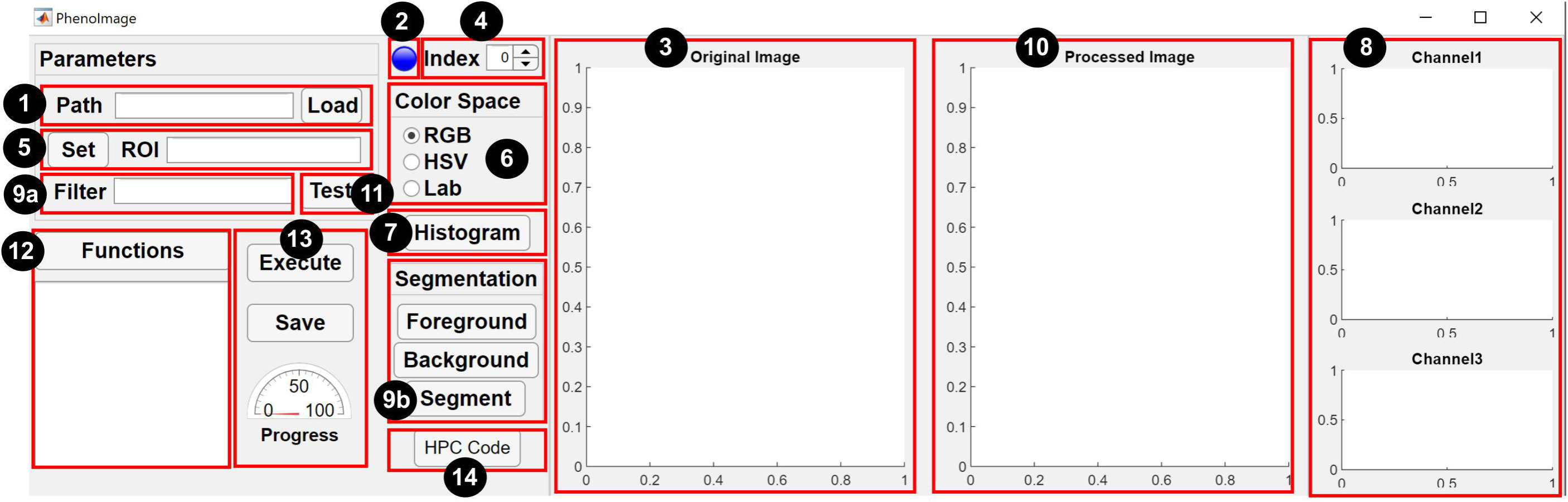
Graphical user interface (GUI) of *PhenoImage*. The numbers denote a step-by-step guide to use the application: (1) define ‘*Path*’ of the folder containing plant images and click ‘*Load*’ button, (2) ‘*Light bulb*’ shows status of the loading procedure, as the blue ‘*Light bulb*’ turns red while the loading is in progress and green when completed, (3) one of the image from the folder is displayed in the ‘*Original Image*’ section, (4) the spinner can be used to change the current image shown in the ‘*Original Image*’ space, (5) then, the user must define ‘*Region of interest*’ or ‘*ROI*’ by dragging the cursor on the *Original image*, (6) select the ‘C*olor space*’ of preference, (7 and 8) click on ‘*Histogram*’ button to visualize intensity of channels corresponding to the respective ‘C*olor space’*, (9a and b) the user can either directly use the histogram values to define the ‘*Filter*’ or the user can use the ‘*Segmentation*’ function, where ‘*Foreground*’ and ‘*Background*’ need to be specified to precisely segment plant pixels from the background, (10 and 11) the user can click ‘*Test*’ to view the ‘*Processed image*’ and if the user has decided on the ‘*Filter*’ (12) then, selection of ‘*Functions*’ is performed. The functions or the digital parameters that need to be extracted are based on user preference. (13) If running on a single machine, the user can ‘*Execute*’ the function to process all images in the respective folder, and progress of batch processing can be viewed in the ‘*Progress bar*’ and saved or (14) high-performance computing clusters can be used to process the images.

The visible images (RGB) of plants can be obtained using any system such as LemmaTec or using standard digital cameras. If analyzing images using standard digital cameras, the user must ensure constant focal distance to have similar scale for all the images corresponding to the same batch to facilitate precise comparison.

### Selection of Region of Interest (ROI) and Image Filtering

For selecting ROI, the user can either crop the image interactively by dragging a marquee tool over the image or by typing the position of the ROI using this format, [X_min, Y_min, Width, Height], where X_min, Y_min is the coordinate of the upper-left corner, and Width and Height correspond to the size of the ROI. The ROI selected is fixed for the image analysis of the respective folder. Thus, it is recommended that the user selects a relatively larger ROI. This is important especially during the analysis of plant growth dynamics in a temporal manner where plants tend to increase in size.

Next, image segmentation separates plant pixels from the background. For segmentation, a logical expression can be specified in ‘*Filter*’ (Fig. 2). The application supports (1) *r*ed, *g*reen, and *b*lue (*RGB*), (2) *h*ue, *s*aturation, and *v*alue (*HSV*), and (3) *Lab* color spaces, which provide flexibility to the user to optimally segment the RGB images. Any combination of arithmetic and logical operation can be used to segment the plant. In terms of setting the filter, ‘&’ means logical *AND*, ‘|’ means logical *OR*. For example, ‘r<200 & g< 150’ means finding pixels that have red values less than 200 and green values less than 150. The ‘*Processed Image*’ will be displayed after clicking ‘*Test*’ button (Fig. 2).

If the user is unsure about predefining the filter, then the ‘*Segmentation*’ feature can be utilized (Zhu *et al*., 2020). For this, click on ‘*Foreground*’ and a new pop-up window having the original image appears. The user can select the zoom-in option from the task bar menu to enlarge the area of interest (i.e., plant tissue in this case). Once the area is zoomed in, the user can deselect the zoom in option and scribble on the enlarged area of interest with a red mark (Fig. 3). Next, background (i.e., pot, pot stand, plant background, etc.) is to be selected by clicking the ‘*Background*’ button and scribble on the background using a green mark. Afterwards, the user can click the ‘*Segment*’ button to initiate the segmentation or subtraction of plant pixels from the background (Fig. 3). We empirically segment the plant by finding plant pixels where the difference to the mean of selected foreground is less than 60. Implementation of the ‘*Segment*’ function may take a few additional seconds. After segmentation, the ‘*Processed Image*’ will show pixels corresponding only to the plant and the histograms corresponding only to the plant region will be displayed in ‘*Channel 1, 2*, and *3*’. The range of the histogram for each channel can be used to define ‘*Filter*’ parameters. The ‘*Segmentation*’ feature is helpful to define filter parameters in a similar manner for both RGB and fluorescence images; however, histogram values for only the red channel need to be considered for setting the filter in the case of fluorescence images.

**Fig. 3.**
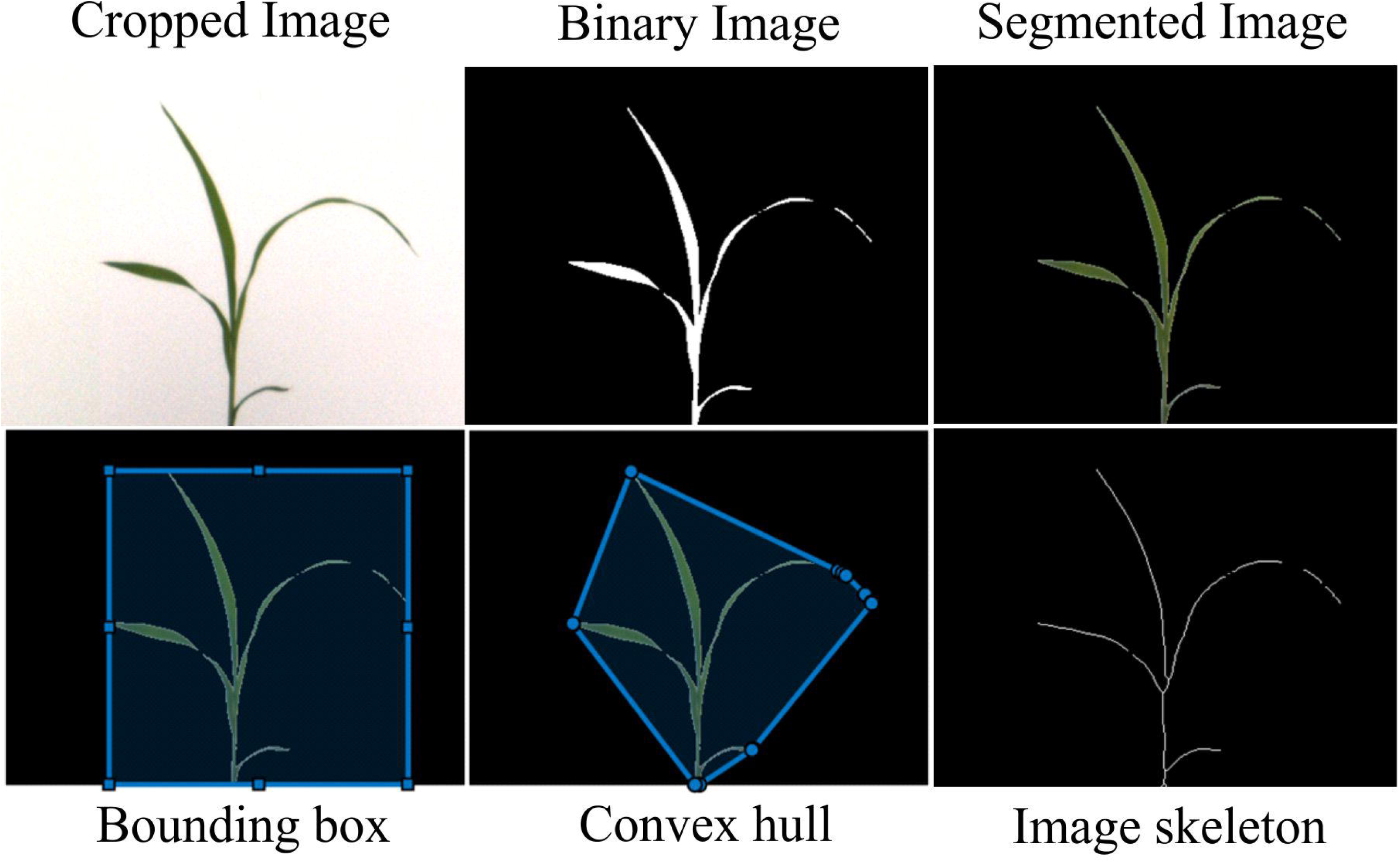
Representation of different features used by *PhenoImage*. The cropped image is derived from the original image after the selection of the region of interest. The binary image is a mask of the plant pixels where the plant pixels are set to 1 and the background is set to 0. The segmented image represents the segmented plant pixels from the background. The bounding box shown in the light blue color is calculated based on the segmented pixels and encloses all pixels of the plant. Convex hull signifies the smallest convex polygon enclosing all the pixels of the plant. The image skeleton approximates the center lines of the stem and the leaves and is calculated using a skeletonization algorithm.

### Defining Functions for Plant Trait Analysis

For digital trait extraction, the user can select functions from a dialog window by clicking ‘*Functions*’, where any user defined functions can be selected. The selected functions will be listed in the text region and will take the segmented image as input to extract digital traits. Some of the commonly used functions are defined below:

#### Pixel Count

After segmentation, only the pixels corresponding to the plant are kept, while pixels corresponding to other objects in the image are set to black. The tool counts the number of pixels that belong to the plant in the ROI.

#### Pixel Intensity

Pixel Intensity refers to the sum of the intensities of pixels in an image. As there are three channels, red, green, and blue, we calculate the pixel sum of each of the R, G, and B channels separately.

#### Dimension

To define the dimensions (width and height), firstly a bounding box, which based on the segmented pixels and encloses all pixels of the plant, is found (Fig. 4). Then, the width and height of the bounding box are used to define the dimensions of the plant.

**Fig. 4.**
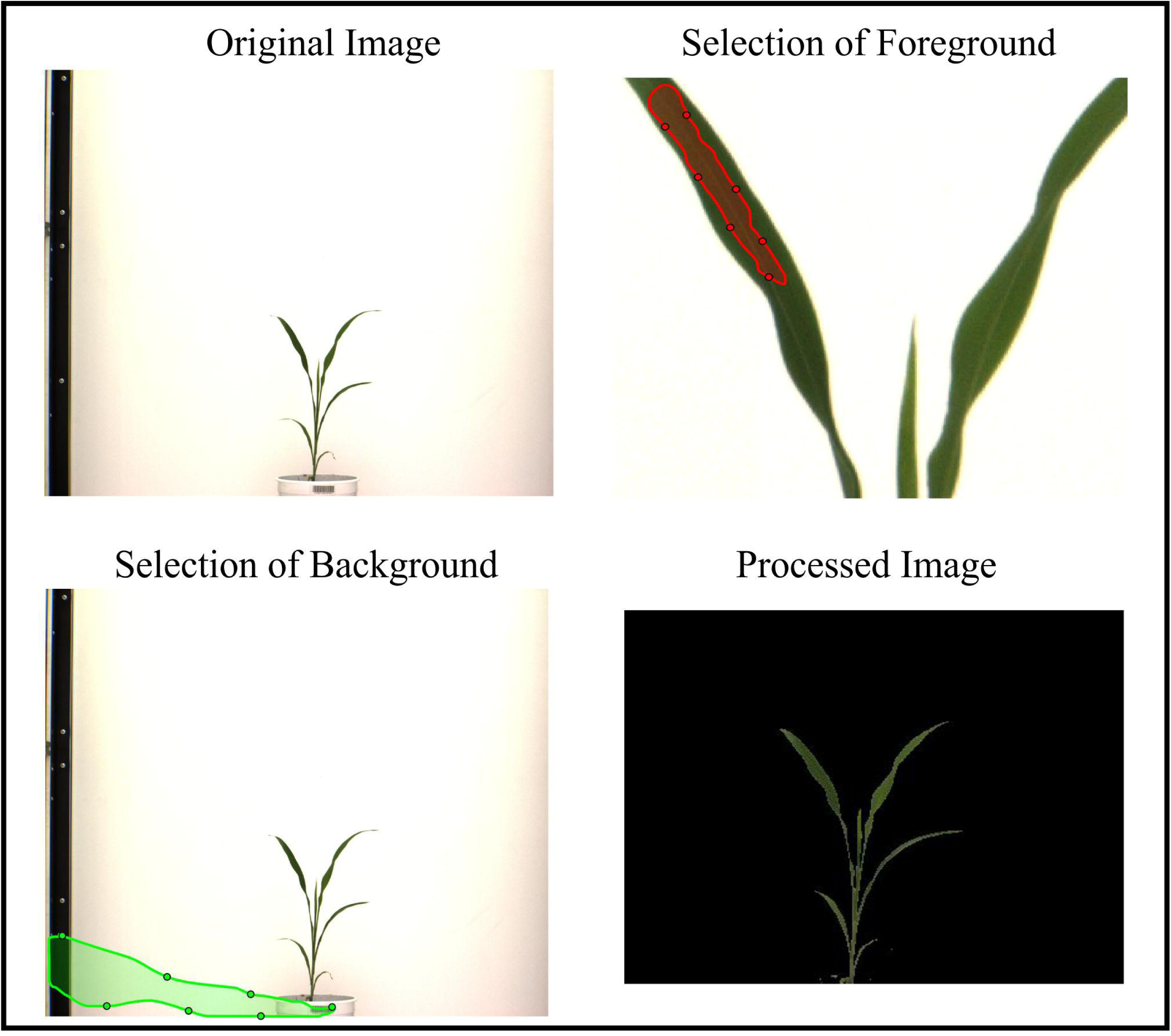
Segmentation. For subtracting plant pixels from the background, click on ‘*Foreground*’ and new a pop-up window displaying the original image opens. The user can scribble on the area of interest with a red mark. Likewise, the user can define background (i.e. pot, pot stand, plant background, etc.) by scribbling on the background using a green mark. Afterwards, the user can click the ‘*Segment*’ button to initiate the segmentation of plant pixels from the background. As a result, the ‘*Processed Image*’ will show pixels corresponding only to the plant.

#### Convex Area

Convex Area is a feature i.e. related to the shape of the plant. The convex area is the area of the convex hull. The convex hull is the smallest convex polygon enclosing all the pixels of the plant (Fig. 4).

#### Image Skeleton

We find the skeleton of the plant pixels using a skeletonization algorithm (Abeysinghe *et al*., 2008). The skeleton of the plant approximates the center lines of the stem and the leaves. Then, the number of pixels in the skeleton is obtained (Fig. 4).

#### Image moment

Image moments can be used to evaluate the shape of the plant (Hu, 1962). We evaluate the image moment of the binary image or the segmented plant image. For the binary image, the plant pixel is considered as 1, and pixels of other objects in the image (e.g., pot, pot-holder, and background) is considered as 0. The moment of the binary image is only dependent on the positions of the pixels. The fourth-order central image moment (μ_22_) is evaluated for the binary image in default. The user can easily modify the function to obtain image moments of other orders.

### Execution on a Local Machine

#### Image Processing and Results collection

The batch image processing can be initiated by clicking ‘*Execute*’ (Fig. 2). The ‘*light bulb*’ located on the right side indicates the status of processing. For instance, the light bulb turns red as images are being processed and will turn green upon its completion. The ‘*Progress*’ gauge will show the progress of the image processing on a percentage basis. A text file containing results from all the functions can be generated by clicking ‘*Save*’. The user can specify the path and file name of the text file in a pop-up dialog window.

### Execution on high performance computing (HPC) clusters

#### Execution and code generation on HPC clusters

For executing jobs on HPC clusters, *slurm* (Yoo *et al*., 2003) is used to submit a batch job, which distributes the jobs using job identifications (IDs). Each job requires a small number of resources, so the priority of execution of each job using *slurm* is high. Thus, owing to less queuing time, we chose *slurm* for *PhenoImage*.

Further, to process images using HPC clusters, click the ‘*Code*’ button (Fig. 2), which generates a MATLAB script for processing images. A *slurm* file (an example is included in *PhenoImage*) is used to submit a job array to the cluster. Then, the job IDs and job size executed by *slurm* are used to partition the images so that each node processes only a part of images. The job IDs and job size are passed to the MATLAB script generated by *PhenoImage* as input parameters. The user needs to input the names of the files that need to be processed in a text file. For each node in the HPC cluster, the script reads all the filenames and processes a part of the images as specified by job ID and job size. Specifically, the script will process images with indices [JobID, JobID+JobSize, JobID+2* JobSize, …]. The script contains the position of the ROI, the expression of the color filter, and the names of the functions that have been selected for digital trait extraction.

After submission of the job using the *slurm* file, the result computed by each function on each node will be printed out and finally aggregated in one output file, which is specified in the *slurm* file. The output file contains all the results for all the input images.

### Sorghum and wheat: a test case for *PhenoImage* validation

#### Sorghum

Four seeds of sorghum genotype, RTX430 were sown in each of the ten 5.6 liter (L) pots (22 cm diameter X 19.5 cm height) filled with 2.5 kg of a soil mix consisting of 2/3 peat moss and 1/3 vermiculite and 1.4 kg lime. Six days after germination, plants were thinned to one seedling per pot. For the first 21 days, all pots were watered to 70% water holding capacity (WHC). Afterwards, water was withheld from half of the pots (water-limited treatment; WL) until 30% WHC is attained, while half of the pots were maintained at 70%WHC (well-watered treatment; WW; (Supplementary Fig. S1). During the entire experiment, the greenhouse was maintained at 28/25°C temperature, 13h/11h – day/night, and 40-50% relative humidity.

#### Wheat

Seeds of wheat genotype – Pavon were germinated in Petri dishes for four days in dark at 25°C. Uniformly germinated seeds were transplanted in 3 L pots (12 cm diameter X 19.5 cm height) filled with 1.2 kg of Fafard germination soil (Sungro, Massachusetts, USA) supplemented with Osmocote fertilizer and Micromax micronutrients. Seedlings were grown for 7 days at 80% WHC. After seven days, 6 seedlings each were maintained at 80% WHC for well-watered treatment and 30% WHC for water-limited treatment (Supplementary Fig. S1). Growth conditions were maintained at 22/16°C – 16/8 h day/night temperatures. Afterwards, plants were imaged every day for 15 days.

#### Water holding capacity (WHC)

For calculating WHC of the soil mix, 2.5 and 1.2 kg of soil mix for sorghum and wheat experiment, respectively, was oven-dried (60°C for 7 days) and dry soil weight was measured. Then, the soil mix was transferred to pots perforated at the bottom for drainage. To achieve the saturation point (weight of the soil at 100% WHC), the soil mix was saturated with water while covered at the top to prevent evaporation. Pots were weighed daily until no change in pot weight was observed. These computed values were then used to calculate the weight of soil at a particular WHC by using the following equation:

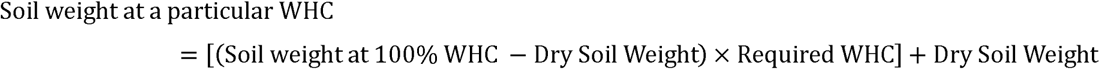

#### Plant Imaging

A high-throughput phenotyping facility (LemnaTec Imaging System) at Nebraska Innovation Campus, the University of Nebraska-Lincoln was used to evaluate sorghum (RTx430) and wheat (Pavon) plants by RGB and fluorescence images. For sorghum plants, starting from the day water was withheld, both WW and WL pots were imaged every day until WL pots reached 30% WHC. Plants were imaged for 18 days (Supplementary Fig. S1). Due to technical error during the experiment, the imaging system failed to acquire images on the 13^th^ day of imaging, so data corresponding to this day is missing from the downstream analysis. For wheat plants, imaging was performed for 15 days for WW and WL conditions (Supplementary Fig. S1).

To reduce image occlusions, imaging was done from five different angles (side views at 0°, 72°, 144°, 216°, and 288°; Supplementary Fig. S2) (Golzarian *et al*., 2011). Next, RGB and fluorescence images from both the species were used as a test cases to validate *PhenoImage*. For validation and optimal segmentation of RGB images, the following filter parameters were used: g<150 (for sorghum) and g<150 & h>0.2 & h<0.5 & s>0.1 & v<0.6 & a<-5 (for wheat).

For fluorescence images, plants were imaged in a separate chamber and an ad hoc-image segmentation strategy was used to categorize image color ranges into 32 color classes (Campbell *et al*., 2015). Further, Hierarchical Cluster Analysis (HCA) was performed using wards method (JMP® Pro13) to examine the temporal profile of the color classes with pixel intensities. For fluorescence images, filters were defined using only the red pixels; sorghum – r>150 and wheat – r>50 & r<140.

### Performance testing

To test the performance of *PhenoImage*, we evaluated the time required to process images with respect to individual functions such as convex area and pixel count (Supplementary Table S1). For this, an RGB image from a sorghum plant was analyzed at resolutions ranging from 100×100 to 10,000×10,000 pixels in an incremental manner.

### Manual phenotyping and comparisons with other methods

Manual measurements were performed for both sorghum and wheat plants on the last day of imaging in a destructive manner. For this, fresh and dry shoots were weighed. Shoots were dried for one week in an oven at 60°C and weighed to determine the dry weight. The manually derived traits were correlated with digital traits derived from last day of imaging for sorghum and wheat (day 18 and 15, respectively; Supplementary Fig. 1). The RGB images from the last day of imaging were processed using *PhenoImage* (Supplementary Table S2) as well as HTPheno (Hartmann *et al*., 2011) and OpenCV (Bradski, 2000) (Supplementary Table S3). For correlation, pixel count derived from *PhenoImage* and OpenCV, and object area from HTPheno were considered.

## Results and Discussion

### Performance Testing

We evaluated the performance of *PhenoImage* with respect to the time required to process images. For this, we computed time required to generate data for two functions: convex area and pixel count, derived from RGB images, which had different levels of resolution ranging from 100×100 to 10,000×10,000 pixels (Supplementary Table S1). We observed that the application’s performance at different resolutions depended on the function that is being evaluated (Fig. 5). For example, time taken to analyze convex area at the highest resolution (10,000×10,000 pixels; 1.646 sec) increased by 53.15% compared to the lowest resolution (100×100 pixels; 0.030 sec). On the other hand, time taken to analyze pixel count increased by 16.20% with the increase in the resolution i.e. 0.167 and 0.009 sec for the highest and lowest resolution, respectively (Fig. 5, Supplementary Table S1).

**Fig. 5.**
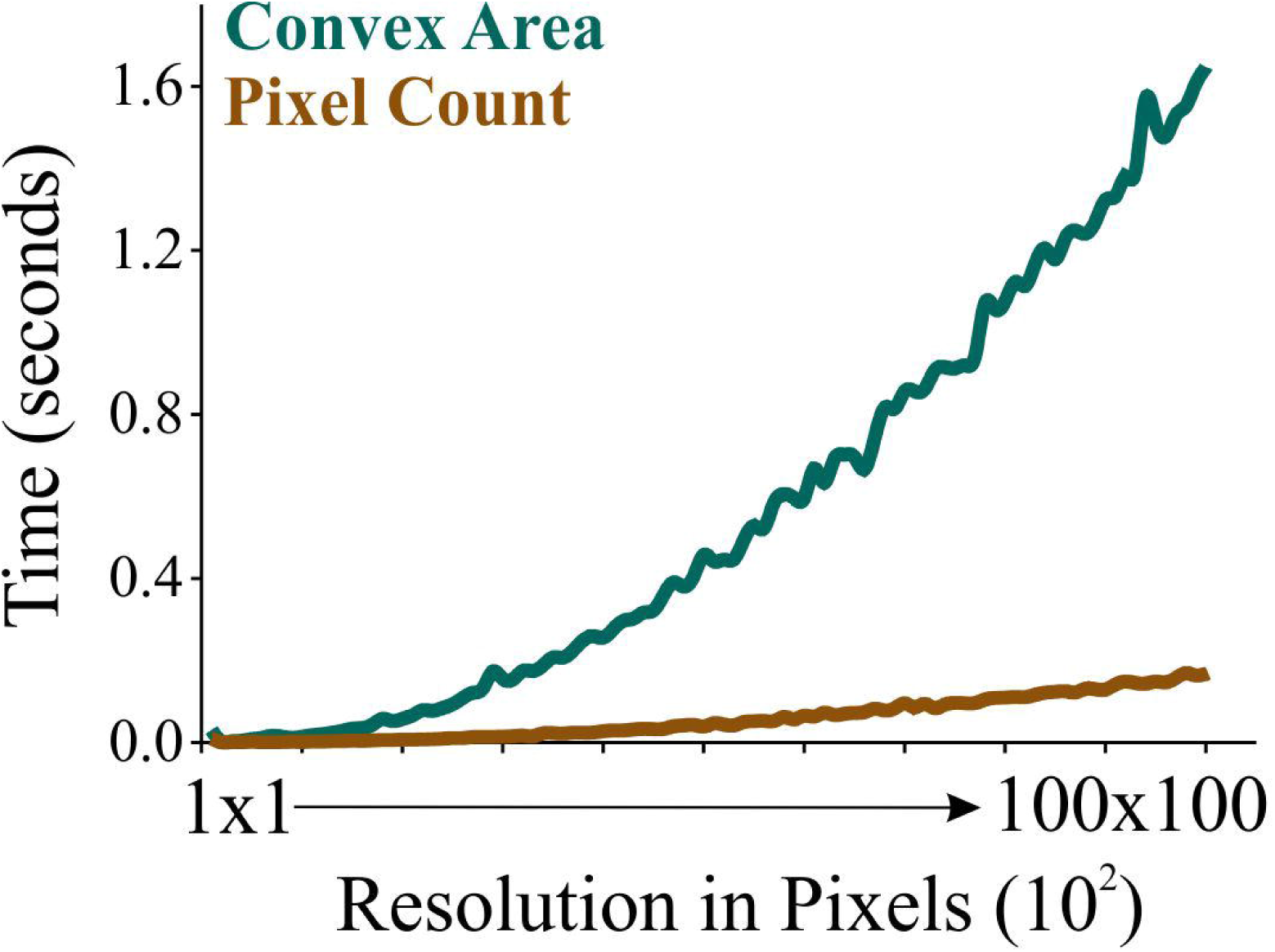
Performance testing of *PhenoImage*. The plot shows the time taken to process images and extract the respective digital trait (convex area and pixel count) from the images at different resolution.

### Sorghum and wheat: A test case for *PhenoImage* validation

The RGB images from sorghum (RTx430) and wheat (Pavon) were used for validating *PhenoImage*. These species were selected because of their visibly different physical attributes. Sorghum has one main shoot axis, which results in a relatively compact-looking phenotype compared to the wheat plant, which produces multiple tillers (Fig. 5). The two species also differ in other parameters such as stalk diameter, leaf width, leaf length, etc. Plants from both the species were used for HTP with the LemnaTec Imaging System, and images were processed using *PhenoImage*. After loading the images onto the application, the best filter parameters were determined empirically based on histogram generated after segmenting the foreground (i.e. plant pixels) from the background.

We assessed two digital traits derived from the RGB images: pixel count and convex area, which are representative of a plant’s overall architecture. Pixel count was used as a proxy for projected shoot area (PSA) and represents the total number of pixels of a plant, whereas the convex area was the area of the convex hull and illustrates the smallest convex polygon enclosing all of the pixels of the plant (Fig. 3). For validation, the visible differences between the two species were assessed for plants of the same age (26-day old) via imaging. As a result, we detected significant differences (*P* < 0.001) between sorghum and wheat plants with respect to PSA as well as the convex area (Fig. 6). Interestingly, although wheat plants have a higher number of leaves than sorghum of the same age, sorghum plants had higher PSA and convex area, apparently due to the broader leaves of sorghum.

**Fig. 6.**
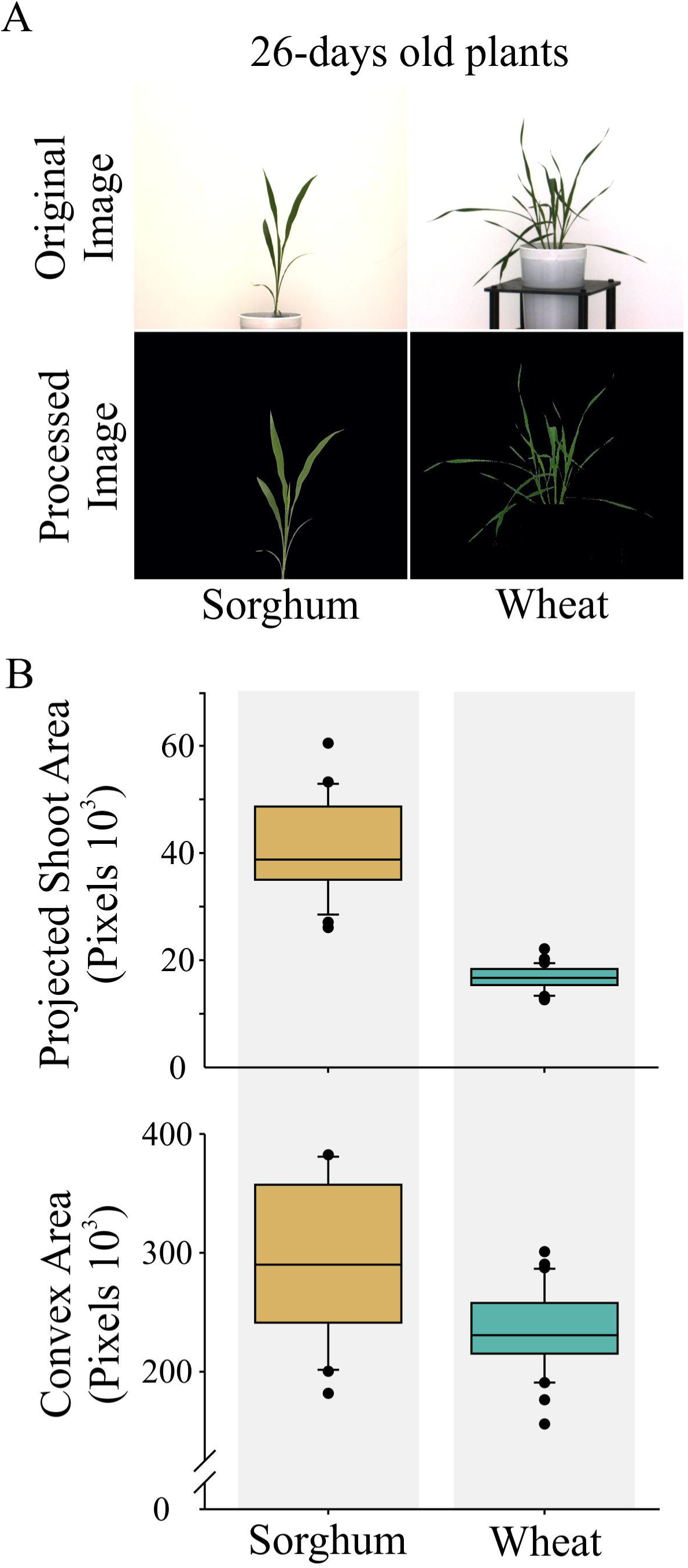
*PhenoImage* validation. (A) The RGB images from 26-day old sorghum and wheat plants. Two digital traits: convex area and projected shoot area (PSA), which represent a plant’s architecture were derived using *PhenoImage*. Significant differences (*P* < 0.001) were detected between sorghum (*n* = 5) and wheat (*n* = 6) for both the digital traits. For statistics, Welch’s t-test (equal variance not assumed) was used.

### Comparison with manual measurements and other image processing methodologies

Next, we performed destructive phenotyping of sorghum and wheat plants at the last day of imaging (day 18 and 15, respectively; Supplementary Fig. S1). The harvested plants were used to manually record fresh and dry shoot weight. As expected, we observed significantly higher fresh and dry weight for sorghum compared to wheat (Supplementary Table S2).

Furthermore, we compared the manually recorded phenotypes with digital traits (pixel count) derived from RGB images. For this, the RGB images were processed using the in-house generated application – *PhenoImage*, as well as two publicly available tools, HTPheno and OpenCV. HTPheno is used as a plugin for ImageJ and does not involve programming language (Hartmann *et al*., 2011). The application does not allow calibration of color settings for image processing. On the other hand, OpenCV requires skills in Python programming language and does not offer GUI (Bradski, 2000). For both the plant species, we detected a high correlation for fresh and dry weight with PSA derived from RGB images processed using *PhenoImage*, which was comparable to correlations obtained from HTPheno and OpenCV (Table 1 and Supplementary Table S2 and S3). This illustrates the sensitivity of the *PhenoImage* application in terms of estimating digital traits from the two plant species while considering minute details in depth.

**Table 1:** Correlation of manual traits – fresh weight (FW) and dry weight (DW) with the digital trait derived

|  | *PhenoImage* | HTPheno | OpenCV |
| --- | --- | --- | --- |
| FW (Sorghum) | 0.923 | 0.897 | 0.901 |
| FW (Wheat) | 0.927 | 0.999 | 0.901 |
| DW (Sorghum) | 0.974 | 0.960 | 0.962 |
| DW (Wheat) | 0.863 | 0.871 | 0.999 |

For comparisons, pixel count derived from *PhenoImage* and OpenCV, and object area from HTPheno for both plant species corresponding to the last day of imaging were used.

### Temporal analysis of growth dynamics using *PhenoImage*

The image-based phenotyping platforms have enabled quantification of physiological and morphological features in a time-dependent manner. In this context, we performed temporal evaluation of sorghum and wheat growth dynamics under well-watered (WW) and water-limited (WL) conditions using HTP. The RGB and fluorescent derived images were processed using *PhenoImage* for testing sensitivity of the tool to detect subtle physiological changes over time.

The biomass of the plant increases with growth and development, which can be quantified by imaging, and environmental stresses in general slow growth and development (Chen *et al*., 2014; Röth *et al*., 2016). To evaluate the changes in plant size in a temporal manner, we traced PSA derived from RGB images under WW and WL conditions. For both sorghum and wheat, PSA showed a gradual increase over time under WW and WL; however, WL conditions exhibited lower PSA relative to WW conditions for the identical time-point (Fig. 7).

**Fig. 7.**
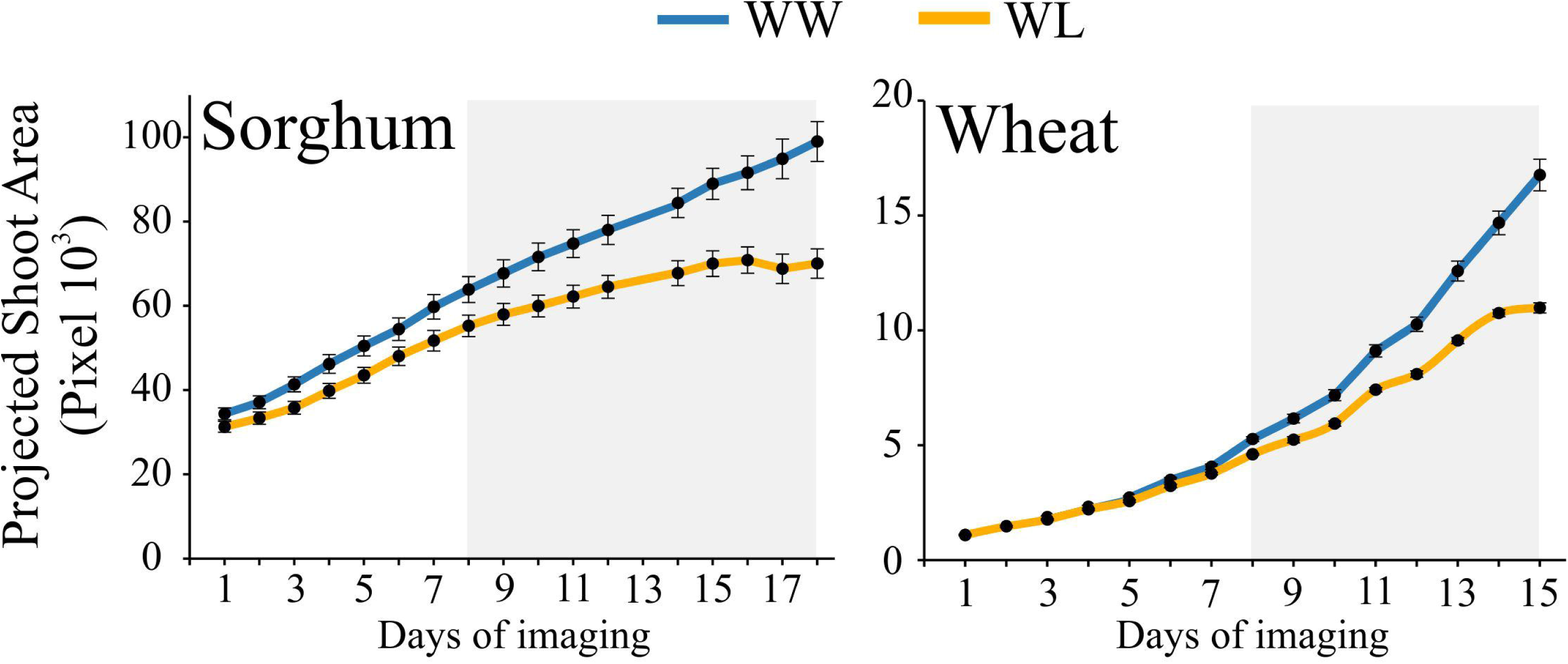
Temporal analysis of growth dynamics. Sorghum and wheat plants subjected to well-watered (WW) and water-limited (WL) conditions were imaged in a time-dependent manner using the LemnaTec Imaging System. Sorghum and wheat plants were imaged for 18 and 15 days, respectively. *PhenoImage* derived projected shoot area (PSA) showed significant differences between WW and WL conditions on the 8^th^ day for both sorghum (*n* = 5) and wheat (*n* = 6). For statistics, the paired *t*-test was used. The grey box represents the significance difference between WW and WL treatments for the respective days.

We evaluated changes in pixel intensities corresponding to the ‘G’ channel and chlorophyll florescence as an indicator of plant health. In principle, the ‘G’ pixel intensity derived from the RGB images reflect the greenness of the plant; higher green pixel intensity reflects higher chlorophyll content, which in turn is associated with the higher photosynthetic activity (Wood *et al*., 2020). The greenness index (GI) was calculated using the following formula: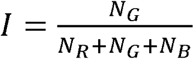, where N_R_, N_G_, and N_B_ are pixel intensity for R, G, and B channel normalized to total pixel count for the respective time-point and treatment. For both sorghum and wheat, we observed higher GI under WW relative to WL conditions (Fig. 8).

**Fig. 8.**
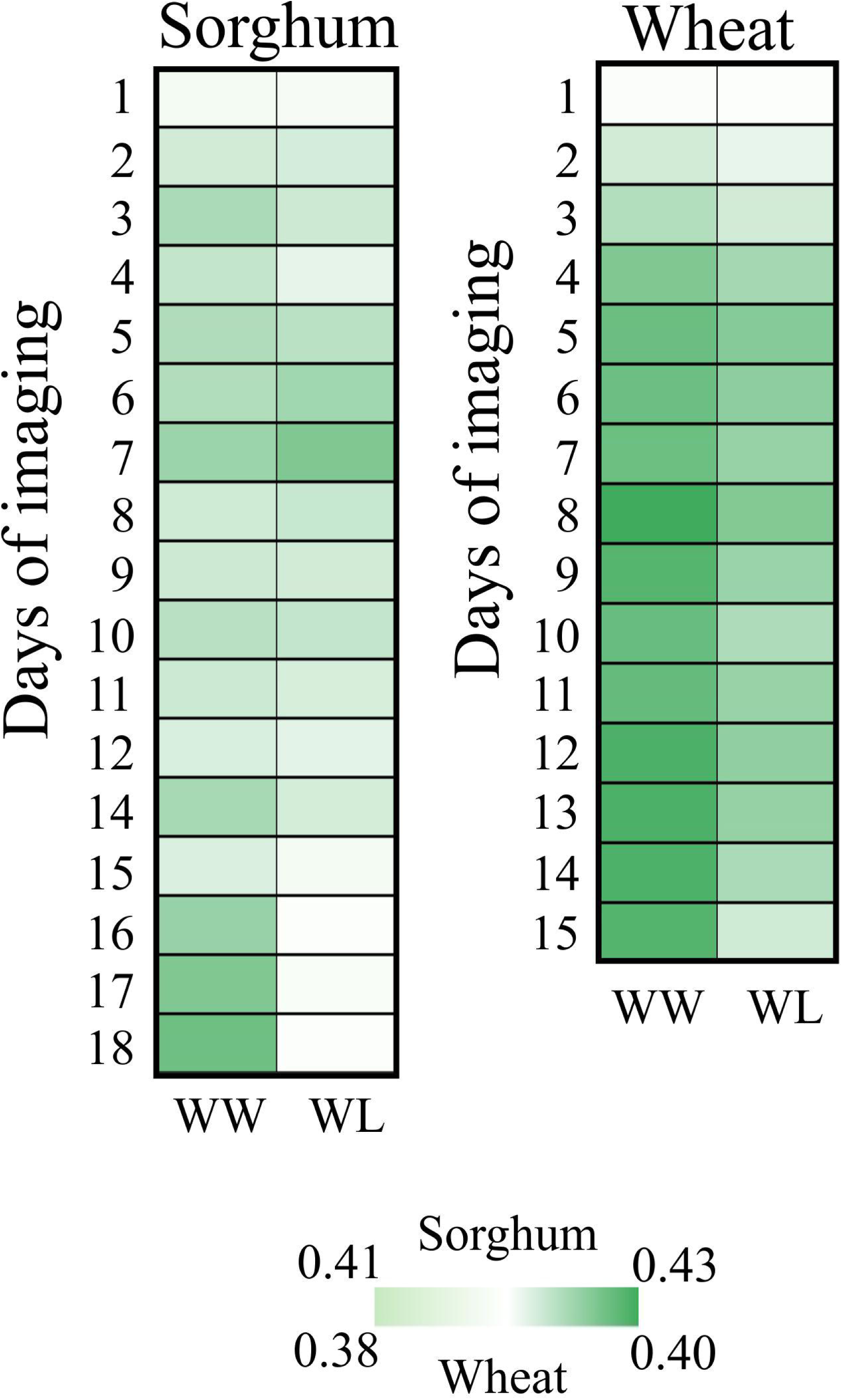
The heat map exhibits the ‘Greenness Index’ for sorghum and wheat under well-watered (WW) and water-limited (WL) conditions in a time-dependent manner.

Furthermore, abiotic stresses such as heat stress or water limitation decreases photosynthetic efficiency and increases non-photochemical quenching resulting in enhanced chlorophyll fluorescence and heat dissipation (Zhao *et al*., 2017; Paul *et al*., 2020). Therefore, we evaluated the dynamics of chlorophyll fluorescence for sorghum and wheat under WW and WL conditions. For this, total pixels corresponding to the red channel were classified into 32 color classes based on their fluorescence intensity. As the stress progressed, fluorescent intensity of pixels changed. To monitor the rearrangement of pixels over time and treatments, we performed HCA. As a result, we detected four clusters (I-IV) each for sorghum and wheat (Fig. 9; left panel). For sorghum, the identified clusters distinguished changes related to both development as well as water treatments (WW and WL). Cluster I comprised fluorescence changes at early time points – 1 to 5 day (d) of imaging, wherein cluster II and III were associated with later time points (d6 to d17) under both WW and WL conditions (Fig. 9). Furthermore, HCA clearly distinguished fluorescence changes linked with water treatment, as cluster II and III were predominant ones under WL and WW conditions, respectively (Fig. 9). In the case of wheat, HCA distinguished development-driven fluorescence changes, as early time points (d1 to d5) were represented by cluster I and II and late time points (d6 to d15) were represented by cluster III and IV (Fig. 9; right panel). However, a clear distinction between WW and WL conditions was not observed. These results are in line with previous findings documenting decreased chlorophyll content or photosynthetic activity as a possible penalty on a plant subjected to WL conditions (Mathobo *et al*., 2017).

**Fig. 9.**
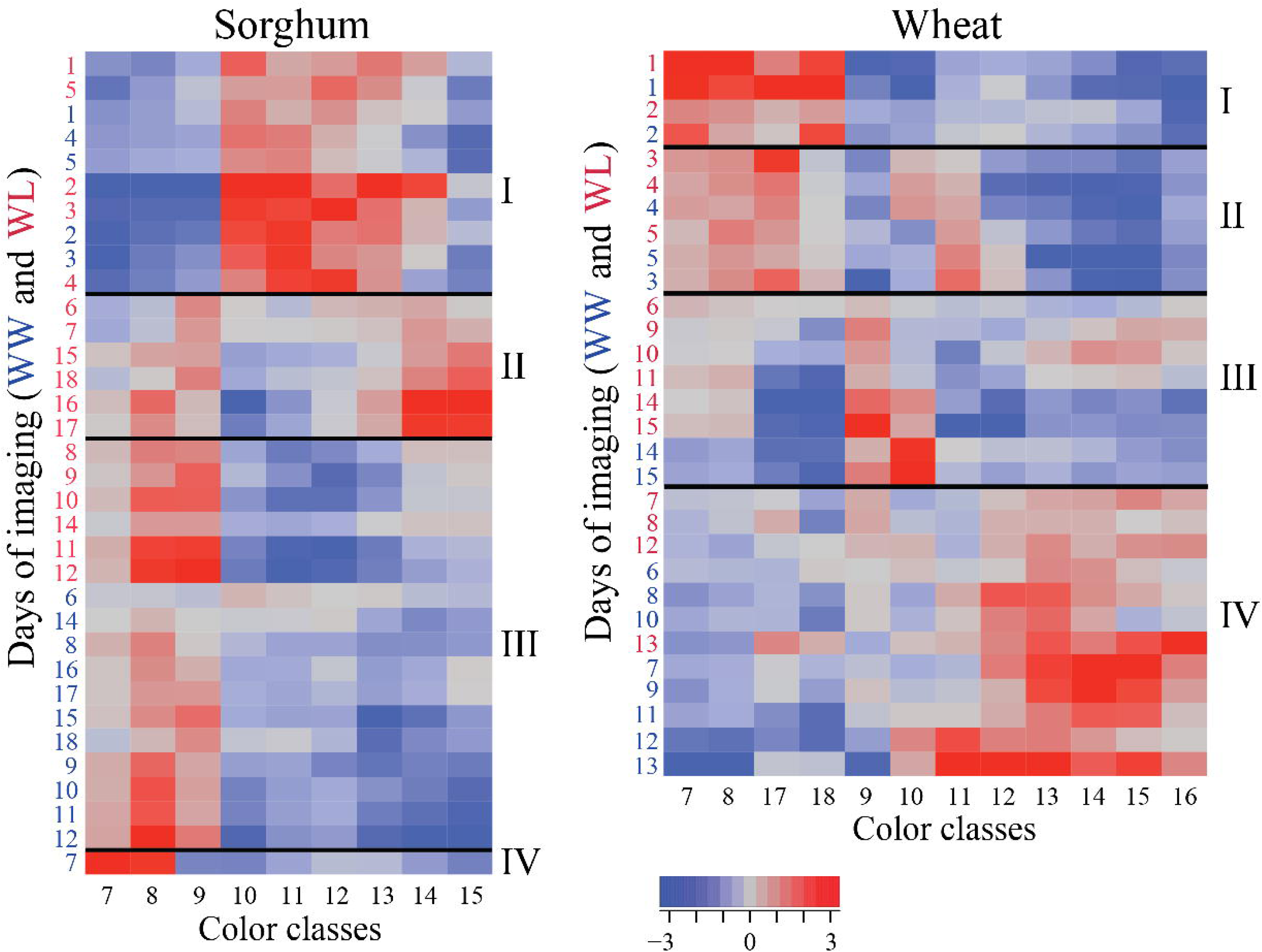
Hierarchical Cluster Analysis of fluorescence color classes for sorghum (left panel) and wheat (right panel). Normalized pixel counts corresponding to different color classes were clustered (I to IV) using wards method in JMP® Pro13 under well-watered (WW) and water-limited (WL) conditions. Days of imaging under WW and WL treated plants were represented by blue and red-colored numerals, respectively.

Collectively, the results establish that *PhenoImage* can be used to analyze HTP-derived longitudinal phenotypic datasets (RGB and fluorescence images) to detect the occurrence of subtle phenotypic changes in a plant’s growth and development.

## Conclusion

*PhenoImage* offers an exhaustive and robust analysis of large-scale plant phenotyping data. The intuitiveness of the application allows scientists with little programming experience to process large-scale datasets on their computers. The tool can also support parallel high performance computing clusters. The availability of multiple functions and filtering parameters provides flexibility to analyze a wide variety of plant species. Moreover, open-source nature provides the possibility to extend the usability of the tool to meet specific user requirements. The current version of the application is designed for analyzing aboveground plant images. However, we plan to extend its usability to examine other tissues such as root or panicles.

## Author Contributions

HW, HY, SS, and PS supervised the project. PP led the study. FZ designed and developed the application. MS and JS performed the experiment. MS, JS, and PP analyzed the results. PP wrote the manuscript with inputs from MS, JS, and FZ. All authors read and approve the manuscript.

## Acknowledgements

We would like to thank the Nebraska Innovation Campus, University of Nebraska, Lincoln, USA for their support for the imaging experiments.

## Software Availability

*PhenoImage* is available in two different versions: (i) Standalone application: this version does not require MATLAB license for its operation, (ii) Regular application: this version does require MATLAB: https://www.mathworks.com/products/matlab.html.

Both versions are available at http://wrchr.org/phenolib/phenoimage. We have provided a detailed step-by-step guide for using *PhenoImage* (*PhenoImage* Guide Document).

## Supplementary Material

Supplementary Fig. S1: Water treatments for sorghum and wheat. For sorghum, all pots were watered to 70% water holding capacity (WHC) for the first 21 days. Then, water was withheld from half of the pots (water-limited treatment; WL) until 30% WHC is attained, while half of the pots were maintained at 70%WHC (well-watered treatment; WW). During the entire experiment, the greenhouse was maintained at 28/25°C temperature, 13h/11h – day/night, and 40-50% relative humidity. For wheat, seedlings were grown for 7 days at 80% WHC. After seven days, half of the seedlings were maintained at 80% WHC for WW treatment. For the other half, water was withheld until 30% WHC is attained (WL treatment). Growth conditions were maintained at 22/16°C – 16/8 h day/night temperatures. Afterwards, plants were imaged every day for 15 days.

Supplementary Fig. S2: Representative RGB original and processed images of sorghum and wheat plants from five different angles.

Supplementary Table S1: To test the performance of *PhenoImage*, we evaluated the time required to process images with respect to individual functions: convex area and pixel count. For this, an RGB image from a sorghum plant was analyzed at resolutions ranging from 100×100 to 10,000×10,000 pixels in an incremental manner.

Supplementary Table S2: Sorghum and wheat plants from the last day of imaging (day 18 and 15, repsectively) were harvested for recording manual traits (fresh and dry weight; FW and DW). These manual traits were correlated with digital traits derived from RGB images processed using PhenoImage. *n* = 5 and 6 for sorghum and wheat, respectively.

Supplementary Table S3: Sorghum and wheat plants from the last day of imaging (day 18 and 15, repsectively) were harvested for recording manual traits (fresh and dry weight; FW and DW). These manual traits were correlated with digital traits derived from RGB images processed using PhenoImage and OpenCV (Pixel count), and HTPheno (Object Area).

## References

Abeysinghe SS, Baker M, Chiu W, Ju T. 2008. Segmentation-free skeletonization of grayscale volumes for shape understanding. IEEE International Conference on Shape Modeling and Applications 2008, Proceedings, SMI.63–71.

Bradski G. 2000. The OpenCV Library. Doctor Dobbs Journal 25, 120–126.

Campbell MT, Knecht AC, Berger B, Brien CJ, Wang D, Walia H. 2015. Integrating image-based phenomics and association analysis to dissect the genetic architecture of temporal salinity responses in rice. Plant Physiology 168, 1476–1489.

Chen D, Neumann K, Friedel S, Kilian B, Chen M, Altmann T, Klukas C. 2014. Dissecting the phenotypic components of crop plant growthand drought responses based on high-throughput image analysis w open. Plant Cell 26, 4636–4655.

Fahlgren N, Gehan MA, Baxter I. 2015. Lights, camera, action: high-throughput plant phenotyping is ready for a close-up. Current Opinion in Plant Biology 24, 93–99.

Fiorani F, Schurr U. 2013. Future Scenarios for Plant Phenotyping. Annual Review of Plant Biology 64, 267–291.

Furbank RT, Tester M. 2011. Phenomics - technologies to relieve the phenotyping bottleneck. Trends in Plant Science 16, 635–644.

Gehan MA, Fahlgren N, Abbasi A, et al. 2017. PlantCV v2: Image analysis software for high-throughput plant phenotyping. PeerJ 2017, 1–23.

Golzarian MR, Frick RA, Rajendran K, Berger B, Roy S, Tester M, Lun DS. 2011. Accurate inference of shoot biomass from high-throughput images of cereal plants. Plant Methods 7, 2.

Gong P, He C. 2014. Uncovering Divergence of Rice Exon Junction Complex Core Heterodimer Gene Duplication Reveals Their Essential Role in Growth, Development, and Reproduction. Plant Physiology 165, 1047–1061.

Hartmann A, Czauderna T, Hoffmann R, Stein N, Schreiber F. 2011. HTPheno: An image analysis pipeline for high-throughput plant phenotyping. BMC Bioinformatics 12, 148.

Houle D, Govindaraju DR, Omholt S. 2010. Phenomics: The next challenge. Nature Reviews Genetics 11, 855–866.

Hu MK. 1962. Visual Pattern Recognition by Moment Invariants. IRE Transactions on Information Theory 8, 179–187.

Jackson SA, Iwata A, Lee S-H, Schmutz J, Shoemaker R. 2011. Sequencing crop genomes: approaches and applications. New Phytologist 191, 915–925.

Klukas C, Chen D, Pape JM. 2014. Integrated analysis platform: An open-source information system for high-throughput plant phenotyping. Plant Physiology 165, 506–518.

Knecht AC, Campbell MT, Caprez A, Swanson DR, Walia H. 2016. Image Harvest: An open-source platform for high-throughput plant image processing and analysis. Journal of Experimental Botany 67, 3587–3599.

Li L, Zhang Q, Huang D, Li L, Zhang Q, Huang D. 2014. A review of imaging techniques for plant phenotyping. Sensors 14, 20078–20111.

Lobet G, Draye X, Périlleux C. 2013. An online database for plant image analysis software tools. Plant Methods 9, 38

Mathobo R, Marais D, Steyn JM. 2017. The effect of drought stress on yield, leaf gaseous exchange and chlorophyll fluorescence of dry beans (Phaseolus vulgaris L.). Agricultural Water Management 180, 118–125.

Minervini M, Scharr H, Tsaftaris SA. 2015. Image analysis: The new bottleneck in plant

phenotyping [applications corner]. IEEE Signal Processing Magazine 32, 126–131.

Moose SP, Mumm RH. 2008. Molecular plant breeding as the foundation for 21st century crop improvement. Plant Physiology 147, 969–977.

Paul P, Mesihovic A, Chaturvedi P, Ghatak A, Weckwerth W, Böhmer M, Schleiff E. 2020. Structural and functional heat stress responses of chloroplasts of Arabidopsis thaliana. Genes 11, 1–20.

Röth S, Paul P, Fragkostefanakis S. 2016. Plant heat stress response and thermotolerance. In: Jaiwal P., Singh R., Dhankher O. (eds) Genetic Manipulation in Plants for Mitigation of Climate Change. Springer, New Delhi.

Tester M, Langridge P. 2010. Breeding technologies to increase crop production in a changing world. Science 327, 818–822.

Varshney RK, Nayak SN, May GD, Jackson SA. 2009. Next-generation sequencing technologies and their implications for crop genetics and breeding. Trends in Biotechnology 27, 522–530.

Wood NJ, Baker A, Quinnell RJ, Camargo-Valero MA. 2020. A Simple and Non-destructive Method for Chlorophyll Quantification of Chlamydomonas Cultures Using Digital Image Analysis. Frontiers in Bioengineering and Biotechnology 8.746.

Yang W, Feng H, Zhang X, Zhang J, Doonan J. 2020. Crop phenomics and high-throughput phenotyping: past decades, current challenges, and future perspectives. Molecular Plant 13, 187–214.

Yoo AB, Jette MA, Grondona M. 2003. SLURM: Simple Linux Utility for Resource Management. Lecture Notes in Computer Science (including subseries Lecture Notes in Artificial Intelligence and Lecture Notes in Bioinformatics) 2862, 44–60.

Zhao X, Chen T, Feng B, Zhang C, Peng S, Zhang X, Fu G, Tao L. 2017. Non-photochemical quenching plays a key role in light acclimation of rice plants differing in leaf color. Frontiers in Plant Science 7, 1968.

Zhu F, Paul P, Hussain W, Wallman K, Dhatt BK, Irvin L, Morota G, Yu H, Walia H. 2020. SeedExtractor: an open-source GUI for seed image analysis. bioRxiv, 2020.06.28.176230.

